# Spatial-temporal order-disorder transition in angiogenic NOTCH signaling controls cell fate specification

**DOI:** 10.1101/2022.12.07.519534

**Authors:** Tae-Yun Kang, Federico Bocci, Qing Nie, José Nelson Onuchic, Andre Levchenko

**Affiliations:** Department of Biomedical Engineering, Yale University; NSF-Simons Center for Multiscale Cell Fate Research, University of California Irvine; Department of Mathematics, University of California Irvine; Center for Theoretical Biological Physics, Rice University

**Keywords:** Angiogenesis, NOTCH Signaling, Order-disorder transition, Tip-Stalk fate, Turing pattern

## Abstract

Angiogenesis is a morphogenic process resulting in the formation of new blood vessels from pre-existing ones, usually in hypoxic micro-environments. The initial steps of angiogenesis depend on robust differentiation of oligopotent endothelial cells into the Tip and Stalk phenotypic cell fates, controlled by NOTCH-dependent cell-cell communication. The dynamics of spatial patterning of this cell fate specification are only partially understood. Here, by combining a controlled experimental angiogenesis model with mathematical and computational analyses, we find that the regular spatial Tip-Stalk cell patterning can undergo an order-disorder transition at a relatively high input level of a pro-angiogenic factor VEGF. The resulting differentiation is robust but temporally unstable for most cells, with only a subset of presumptive Tip cells leading sprout extensions. We further find that sprouts form in a manner maximizing their mutual distance, consistent with a Turing-like model that may depend on local enrichment and depletion of fibronectin. Together, our data suggest that NOTCH signaling mediates a robust way of cell differentiation enabling but not instructing subsequent steps in angiogenic morphogenesis, which may require additional cues and self-organization mechanisms. This analysis can assist in further understanding of cell plasticity underlying angiogenesis and other complex morphogenic processes.

**Significance Statement:** We investigate the spatial and temporal patterns of Tip/Stalk specification and the ensuing angiogenic sprouting by using a novel controlled micro-engineered experimental model of angiogenesis and a set of mathematical models of the spatially resolved, cell population-level VEGF-NOTCH signaling. Our analysis provides a dynamic view of the initial step of angiogenesis, revealing fluctuations in its onset, and features suggesting transitions between order and disorder in cell organization. These findings suggest how a potentially very restrictive patterning mechanism can become sensitive to a variety of environmental cues. This sensitivity can be crucial for proper vascularization of a damaged organ, and may suggest new ways of analyzing angiogenesis in the context of cancer and other pathologies. This analysis also suggests a framework for understanding of other instances of NOTCH-mediated patterning processes.

## Introduction

Angiogenesis, i.e., the formation of new blood vessels from the pre-existing ones, is a striking example of phenotypic plasticity in an adult differentiated endothelium. Pro-angiogenic factors secreted in response to hypoxic conditions, particularly the vascular endothelial growth factor (VEGF), specify differentiation of endothelial cells lining blood vessels into diverse phenotypic states, including the pro-migratory Tip cell phenotype. Tip cells can initiate invasive cell migration into the surrounding extracellular matrix (ECM), leading to sprouting and branching of the nascent vessels(1, 2). Tip cells are differentiated from Stalk cells, another phenotypic state, through juxtacrine cell-cell interaction between these cell types involving NOTCH1 signaling, triggered and modulated by induction of Dll4 and Jag1 ligands(3, 4). Stalk cells can therefore form in immediate proximity of Tip cells, particularly, at the leading edge of an extending sprout, if the NOTCH signaling is sufficiently pronounced for the Tip-Stalk differentiation to occur. Proliferation of Stalk cells is as essential as the invasive migration of Tip cells for the emergence, extension and branching of growing sprouts, making the analysis of coordinated Tip and Stalk specification particularly important.

The inputs specifying the cell fate can be potentially contradictory, e.g., with pro-angiogenic factors, such as VEGF, promoting the Tip cell fate, and the NOTCH signaling activated by the neighboring cells promoting the Stalk cell fate and thus suppressing the Tip cell identity in the same cell. These and other signaling inputs can thus be incoherent in terms of cell fate specification and can result in complex dynamic outcomes that are still poorly understood. Further elucidation of these processes thus requires high resolution, quantitative experimental measurements tightly coupled with computational analysis. Since such measurements are still challenging *in vivo*, particularly in mammalian tissues, use of tissue models recapitulating the salient features of the developing vasculature is a key tool in the current analysis of angiogenesis and development of *de novo* vascular beds.

Previously, we and others have developed a set of micro-fabricated experimental angiogenesis models that have had progressively improved biomimetic characteristics(5-9). These characteristics include spatially and biochemically appropriate cell micro-environments, composed of components of the extracellular matrix and of gradients of growth factors and cytokines around the developing vasculature, which is composed of endothelial cells and pericytes.

We have previously used this approach to map different combinations of VEGF and an inflammatory cytokine, TNF, onto pro- and anti-angiogenic outcomes, modeling frequently encountered angiogenesis conditions(7). This analysis provided evidence that ‘mini-sprouts’ — one-cell structures protruding from the parental blood vessel into the surrounding matrix — were comprised of Tip cells. However, it was not clear whether all such ‘mini-sprouts’ would ultimately develop into more mature multi-cellular sprouts with defined lumens and the potential to form new blood vessels. Furthermore, although our analysis was successful in explaining the fraction of Tip cells formed under different conditions, it was not clear how to account for the spatial aspects of the Tip cell and mature sprout specification, such as their mutual separation and density.

We address these challenges here by extending our analysis to a higher temporal and spatial resolution, both in experimental and mathematical models of angiogenic sprouting. Surprisingly, we found that the formation of mini-sprouts was a highly dynamic process, in which they could either retract after extension or form full-fledged sprouts. Furthermore, the experimentally determined spatial positioning of mini-sprouts was well explained by the predicted locations of the Tip cells in the mathematical model but the model could not account for which of the mini-sprouts would become fully formed sprouts. Further analysis revealed that the stable sprout formation from mini-sprouts can be enabled by the local fluctuations of the density of fibronectin, a key pro-angiogenic ECM component, leading to sparse patterns where sprouts tend to maximally distance themselves from other fully formed sprouts. These results reveal some of the key mechanisms that may define the density of the angiogenically formed vascular beds under diverse conditions.

## Results

### Dynamic angiogenesis can be explored in a 3D biomimetic experimental setup

To investigate the properties of angiogenic patterning and cell fate specification, we used an experimental model previously employed to assess the crosStalk between pro-angiogenic and pro-inflammatory stimuli(7). In this experimental setup, angiogenesis occurs from a 3D parental engineered endothelial vessel embedded in the collagen matrix and exposed to exogenously supplied VEGF and other pro-angiogenic factors (Fig. 1A-B). In agreement with prior observations, we found that this setup resulted in formation of both one-cell extensions into the matrix (mini-sprouts) and full-fledged multi-cellular sprouts containing detectable lumens and pronounced leading Tip cells (Fig. 1C). Sprouts displayed a variety of growth stages, including the very early ones, composed of one lumenized cell or pairs of connected cells, also forming a lumen (Table 1). Although mini-sprouts formed throughout the observation area of the vessel, sprouts developed within specific zones, while other zones remained devoid of detectable sprout formation over the course of the study (Figs. 1D-F). These observations suggested that cell fate specification and sprout formation are dynamic processes that may display diverse local outcomes. We therefore set out to characterize these processes in the context of an accessible and well defined analysis tool that can allow to contrast experimental findings with mathematical models of angiogenic patterning, particularly those based on the commonly assumed NOTCH receptor mediated cell-cell interactions (Fig. 1G).

**Table 1.**
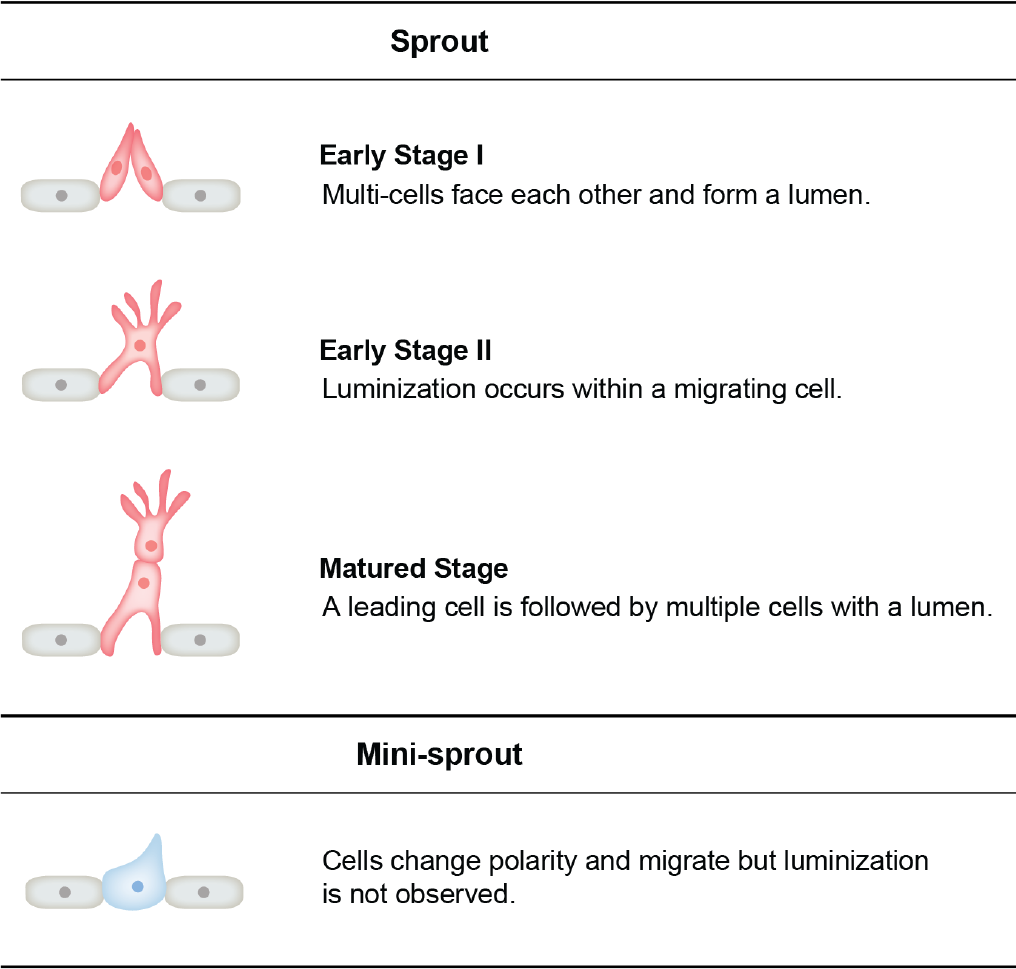
Phenotypic categorization of sprout and mini-sprout.

| Sprout |  |
| --- | --- |
|  | **Early Stage I**<br>Multi-cells face each other and form a lumen. |
|  | **Early Stage II**<br>Luminization occurs within a migrating cell. |
|  | **Matured Stage**<br>A leading cell is followed by multiple cells with a lumen. |
| Mini-sprout |  |
|  | Cells change polarity and migrate but luminization is not observed. |

**Figure 1.**
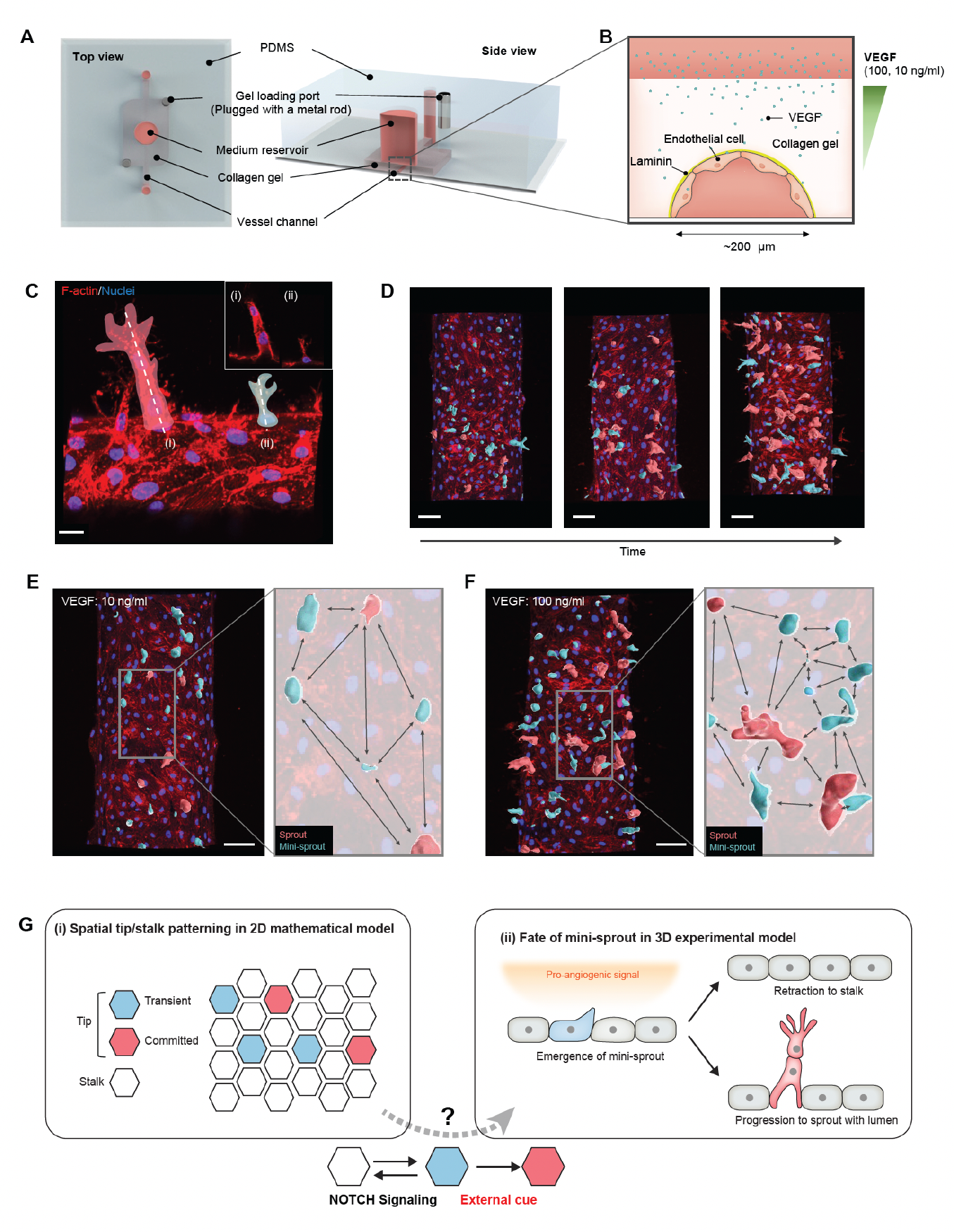
Analysis of temporal and spatial regulation of angiogenic fate specification in a 3D biomimetic experimental setup. (A) 3D vessel model for inducing angiogenesis in response to a gradient of VEGF. (B) Cross-section of a vessel embedded in collagen type I within the device; VEGF is added to the medium reservoir above the vessel to generate a VEGF gradient. (C) Angiogenesis leads to the formation of new sprout-related structures from the parental vessel that have two distinct morphologies: (i) full-fledged multi-cellular sprouts containing detectable lumens, (ii) mini-sprouts in the form of single cell extension into the matrix. Scale bars: 20 *μm*. (D) Temporally resolved observation of dynamic formation of sprouts and mini-sprouts populations during angiogenesis. As depicted in (C), sprouts are pseudo-labeled with red color and mini-sprouts in blue color. Scale bars: 50 *μm*. Dependence of the spatial distribution of sprouts and mini-sprouts on the VEGF concentration: (E) 10 ng/ml and (F) 100 ng/ml. Scale bars: 50 *μm*. Images are 3D reconstructions of confocal z-stacks, showing nuclear (DAPI) and cytoskeleton (Phalloidin). (G) Schematic overview of Tip-Stalk patterning: (i) Spatial Tip-Stalk patterning due to juxtacrine NOTCH signaling that might lead to fixed persistent and transient cell fate specification. (ii) Fates of mini-sprouts in experiments: both retraction (thus conversion from the phenotypically Tip to as Stalk phenotype) and stabilization and growth to a fully defined sprout are observed.

### Mathematical model of VEGF/NOTCH signaling predicts spatially resolved Tip-Stalk patterns

To set the framework for the analysis of cell fate determination, we extended our previously developed and experimentally validated mathematical model of Tip-Stalk fate differentiation between two cells(10) to a multicellular hexagonal lattice in two-dimensions. In the new model, we replicated within each cell the signaling network incorporating the NOTCH and VEGF pathways (Fig. 2A). Prior analysis of this model on the level of two adjacent cells predicted the emergence of bistability between a (high NOTCH, low Delta) Stalk phenotype and a (low NOTCH, high Delta) Tip phenotype(10). This result is consistent with the overall expectation of the differentiation effect of Delta-NOTCH signaling, which in 2D is further expected to generate ‘salt-and-pepper ‘patterns, with a single Tip cell surrounded by 6 Stalk cells (11), yielding the overall fraction of Tip cells in this arrangement of 25%. However, this simple bistability and spatial patterning picture can be altered by signaling inputs that potentially conflict with those involved in Delta-NOTCH signaling (Fig.1G). For example, the VEGF pro-angiogenic factor promoting the Tip cell fate, can conflict with the NOTCH signaling activated by the neighboring cells that instead promotes the Stalk cell fate while suppressing the Tip cell identity. This might result in disordered patterns with adjacent Tip cells that deviate from the archetypical salt-and-pepper configuration. To explore the properties of this disordering effect, we ran simulations, in which the VEGF-NOTCH signaling occurred in all individual cells within a hexagonal array of the model multi-cellular endothelium, starting from randomized initial conditions (Fig. 2A). The fully equilibrated patterns were then analyzed for distributions of the simulated Delta and NOTCH expression across the cells. We found for a wide range of VEGF inputs that the distributions of Delta and NOTCH displayed largely bimodal distributions, and the levels of the average Delta expression increased with the increasing input (Fig. 2B-C, supplementary Fig. 1A-B) due to positive effect of the activated VEGF receptor on Delta (see again the circuit in Fig. 2A). Nevertheless, the clear overall bimodality allowed us to consistently classify cells into the Delta-high (‘Tip’) and Delta-low (‘Stalk’) cell states and examine the spatial distribution of these cellular sub-types (see Method section: Definition of Tip cells in the model).

**Figure 2.**
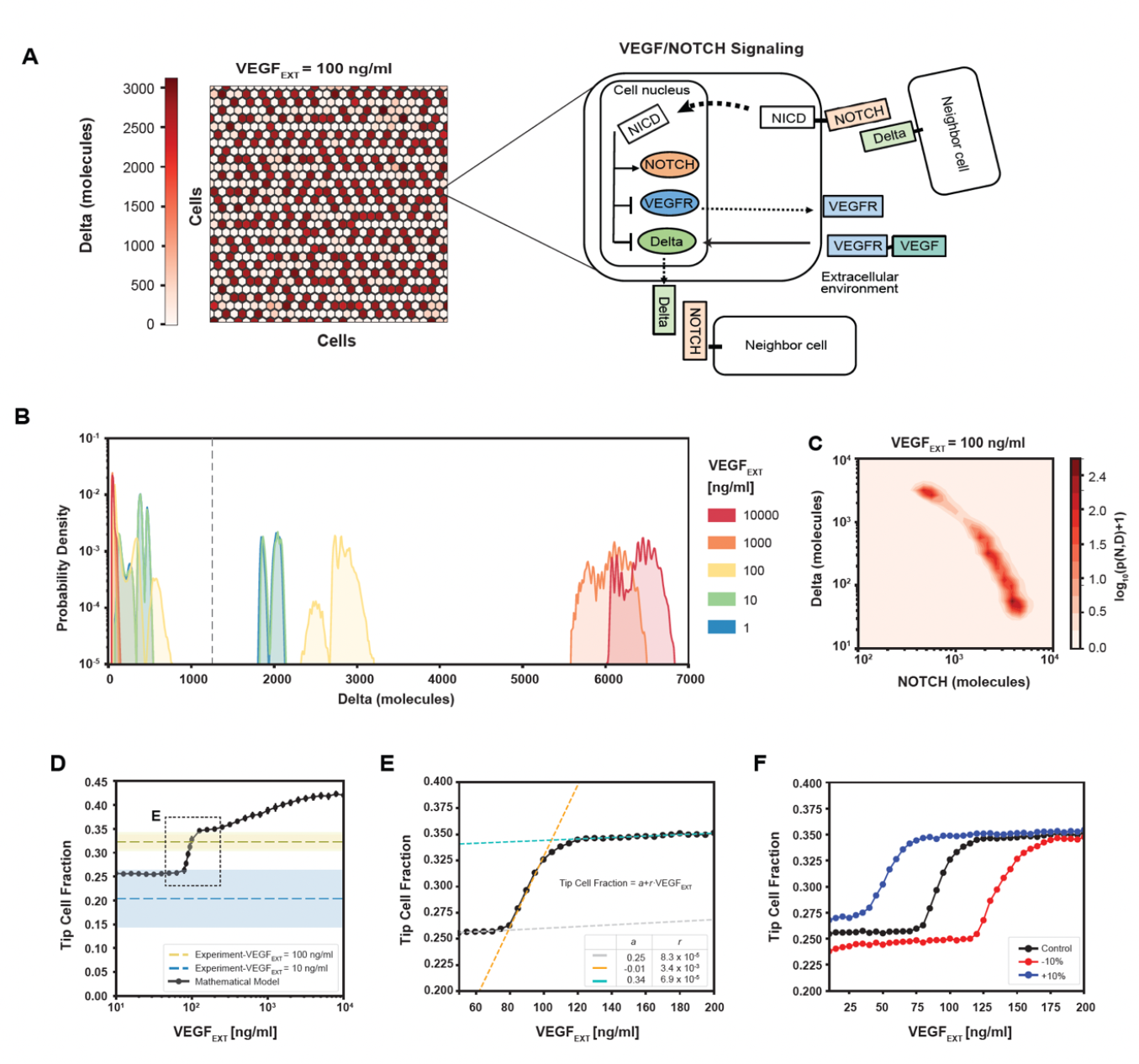
Robust differentiation and order-disorder transition are suggested by mathematical and experimental analyses. (A) Right: An example of a pattern after full equilibration on a 30x30 hexagonal lattice. Color scale highlights the intracellular levels of Delta. Left: The circuit schematic highlights the components of the intracellular NOTCH-VEGF signaling network. (B) Distribution of intracellular Delta levels in the two-dimensional lattice for increasing levels of external VEGF stimuli. (C) Pseudopotential landscape showing the distribution of intracellular levels of NOTCH and Delta for VEGF_EXT_=100 ng/ml. (D) Fraction of Tip cells as a function of external VEGF stimulus (black curve). Blue and yellow lines and shading depict experimental fractions of Tip cells for VEGF_EXT_=10 ng/ml and VEGF_EXT_=100 ng/ml, respectively. (E) Detail of Tip cell fraction transition zone (corresponding to box in panel D). Legend depicts the coefficients of the linear fits. (F) Shift of the VEGF_EXT_ transition threshold upon variation of the NOTCH-Delta binding rate constant. For panels B-F, results are averaged over 50 independent simulations starting from randomized initial conditions for each VEGF_EXT_ level (See Methods section: Simulation details).

The spatial Tip-Stalk cell distribution patterns revealed a complex dependency on the VEGF input. At relatively low VEGF levels, the patterns were mostly ordered, with small deviations from the expected ‘salt and paper ‘geometry with a 25%-75% ratio of Tip-Stalk (Fig. 2D). However, as the VEGF input increased, the fraction of Tips grew and the patterns became sharply more disordered over a relatively narrow range of magnitude of the VEGF input, which could be identified as a highly sensitive area separating more ‘ordered-like’ and ‘disordered-like’ patterns. Finally, increasing VEGF stimuli beyond the highly sensitive area further increased the disorder of the patterns, but with a lower VEGF sensitivity, over several more orders of magnitude of VEGF inputs (Fig. 2D-E and supplementary Fig. 1A-B). Spatial patterns in the disordered phase at high VEGF input levels were characterized by much higher fractions of Tip cells that were frequently in contact with each other as quantified by a “disorder index” (supplementary Fig. 1C-D). This transition was reminiscent of the order-disorder transition commonly observed and studied in the change of the order of atoms in various substances as a function of temperature, and in other chemical systems (12, 13). As expected for these types of transitions, increasing or decreasing the order-stabilizing Delta-NOTCH cell-cell signaling, resulted in the corresponding shifts of the ranges of VEGF inputs, over which the sharp order-disorder transition occurred. For example, a small increment or decrease in the cellular production of the Delta ligand shifted the order-disorder transition to either lower or higher VEGF inputs (Fig. 2F). As the broad sweep of the VEGF inputs simulated in the mathematical model is likely beyond the range of receptor sensitivity for the experimental VEGF signaling, we contrasted the predicted fractions of Tip cells with the previously made observations in the 3D experimental angiogenesis model shown in Fig. 1. We found that, for the VEGF = 10 ng/ml, the experimentally determined fraction of the Tip cells was 0.20 ± 0.08 (mean ± SD), i.e., encompassing the fraction of 0.25 expected for the completely ordered, ‘salt-and-pepper ‘Tip cell distribution pattern. However, for VEGF = 100 ng*/*ml the experimentally observed Tip cell fraction was 0.32 ± 0.01 (mean ± SD), which corresponded to the disordered state predicted by the model under higher VEGF inputs. We therefore concluded that the transition between order and disorder can occur in the 10-100 ng/ml range of VEGF concentration, allowing us to investigate the properties of the disordered state and its relationship to the spatial frequency of sprout formation. Based on the Tip/Stalk cell ratio, we calibrated the model’s parameters so that a VEGF input of 100 ng/ml matched the experimentally observed Tip/Stalk fractions at the same experimental VEGF input (see Methods section: Mathematical model of VEGF/NOTCH signaling).

### Precise quantification of Tip cell spatial arrangement suggests disordered patterning in the engineered angiogenesis model

To enable the comparison between the modeling predictions and experimental observations, we first quantified the spatial patterning characteristics of the inferred positions of the Tip cells in the experimental angiogenesis model. As in our prior analysis using this experimental approach (7), we identified Tip cells based on their key *phenotypic* characteristic — invasive migration into the surrounding collagen matrix. As suggested above, Tip cells can either be present in the form of ‘mini-sprouts’ or be at the Tips of sprouts containing recognizable lumens (see Fig. 3A and Table 1 for examples of this classification). Since formation of both mini-sprouts and lumenized sprouts involved emergence of Tip cells specified in endothelial cell monolayer lining the parental vessel, we constructed a two-dimensional map of the experimentally inferred spatial positions of Tip cells at the location of all mini-sprouts and sprouts (Figs. 3A & B). This mapping (see Methods section: Quantification of Tip-Tip cell distance in experiments) assumed that the Stalk cells found in the extending sprouts emerge through cell proliferation, rather by Stalk cell migration from the parental vessel. Experimentally, we used nuclear staining to identify non-Tip cells. Since the results above suggested that the cell fates and their patterns in our experimental setup were consistent with the ordered or somewhat disordered ‘salt-and-paper ‘patterns, we further assumed that all the non-Tip cells adopted the Stalk fates (in the sense of Delta-low, NOTCH-high status, alternative to the Tip cell fate). The resulting map of experimentally specified Tip and Stalk cell locations was then used to calculate the shortest distances between Tip cells, measured in ‘cell hops’, i.e., the minimal number of intermediate cells between randomly chosen pairs of Tip cells (Fig. 3B). These distances included Stalk cells exclusively, and no intermediate Tip cells. If two or more Tip cells were at equal distance from a given other Tip cell, their distance ranking was assigned randomly (e.g., two Tip cells at the equal minimal distance to a given Tip cell, would be randomly assigned the ranking of the closest and second closest Tip cell). We then analyzed these data for the VEGF_EXT_= 100 ng/ml experimental input which resulted in the most robust sprouting, comparing the results with the predictions of our mathematical model for the same input level that, when averaging over mulTiple simulations, matched the experimentally measured fraction of Tip cells. Finally, we quantified the shortest paths separating Tip cells in the equilibrated patterns (Fig. 3C and Methods section: Quantification of Tip-Tip cell distance in modeling).

**Figure 3.**
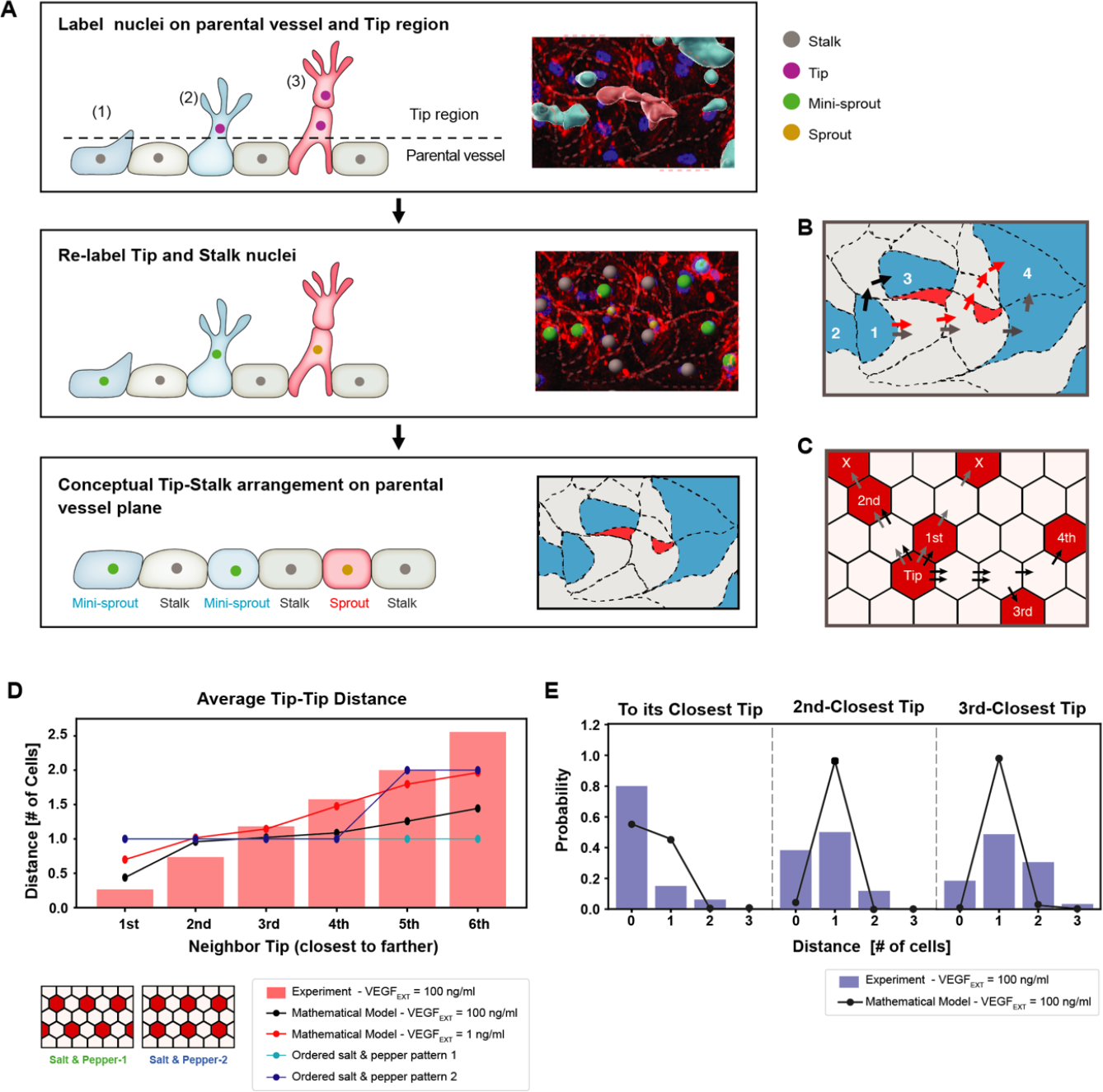
Experimentally measured spatial distribution of Tip cells defined as constituting mini-sprouts and leading sprouts is consistent with the mathematical model predictions. (A) Analysis pipeline to infer the 2D Tip-Stalk arrangements from 3D experimental images: experimental labeling of the nuclei of sprout/mini-sprout cells (above the plane of the parental vessel) and of the Stalk cells (below the plane of the parental vessel) is used to ‘compress’ the cells in each sprout or mini-sprout into a single Tip cell. Tip-Tip distance is defined as the number of cells measured in ‘cell hops’, i.e., the minimal number of intermediate cells between randomly chosen pairs of Tip cells from experiments (B) (the example is identical to the inset at the bottom of (A)) and 2D Tip-Stalk patterns from mathematical modeling (C). Black arrows indicate minimal and valid cell hops between Tips, whereas grey arrows indicate minimal but invalid cell hops (passing through other Tip cells); red arrows indicate the non-minimal cell hops which does not count in the Tip-Tip distance quantification. (D) Tip cell distance distribution from any given Tip cell. Red bars depict experimental measurement for VEGF_EXT_=100 ng/ml and black line depicts the model’s prediction for VEGF_EXT_=100 ng/ml. For reference, dashed lines indicate the expected Tip-Tip distance distribution of “perfect” salt-and-pepper patterns shown in the inserts. (E) Detailed distance distribution for the closest Tip (left), second closest Tip (middle), and third closest Tip (right).

The processing of experimental data described above permitted a direct comparison of modeling predictions and experimental defined Tip cells locations, First, we examined the average distances from the Tip cells to the closest, second closest, etc. neighboring Tip cells. We found that this distance distributions closely agreed with the modeling predictions, particularly for the distances up to the 4th closest neighbor, differing substantially from the predictions for the ordered ‘salt-and-pepper ‘patterns (Fig. 3D). A key finding was that, in agreement with the expectation from the disordered pattern model, there were frequent cases of direct contact between two Tip cells, making the average distance to the closest Tip cell around 0.5 cells in the model. In contrast, two Tip cells were always separated by at least one Stalk cell in “traditional” ‘salt-and-pepper ‘models. As a baseline comparison, the mathematical model with a 100-fold reduction of VEGF stimulus (1 ng/ml) exhibited a Tip-Tip distance statistics more closely comparable with the ‘salt-and-pepper’ models. Further analysis of the experimental distributions of Tip cell distances revealed that Tip cells were adjacent to at least 1 other Tip cell with 80% chance, and with at least 2 other Tip cells with 40% chance, and with at least 3 other Tip cells with 20% chance (Fig. 3E). These Tip cell distance distributions were again in agreement with the modeling results. Taken together these findings provided strong evidence for the predicted partially disordered pattern of Tip cell specification. In our experiments, the observed cell-cell contact area varied, spanning from almost corner-to-corner contact up to approximately 50μm. Previous studies(14, 15) have clearly demonstrated the influence of the cell-cell contact area on NOTCH Signaling, but the values get nosy in the middle range, particularly when excluding extremely low cell-cell contact areas. Reflecting these findings, we excluded the corner contacts, which might correspond to extremely low cell-cell contact areas, from the Tip-Tip distance measurements as depicted in Fig. 3B. We also made an assumption that variations in cell-cell contact size within tens of microns correlate weakly with the strength of NOTCH signaling. This assumption did not impede our effort to compare the overall trends with results from modeling using hexagonal cells, as shown in Figs 3 D&E.

### Dynamic tracking of angiogenic cell fate specification

The integrative, computational and experimental, analysis presented above suggested that the spatial Tip cell distribution can be well explained by the model of a partially disordered ‘salt-and-pepper’ mechanism. However, it is not clear whether all such Tip cells would spearhead the formation of a new sprout, or retract back to an alternative (Stalk) cell fate. We addressed this question by dynamically tracking the fates of mini-sprouts to examine whether this state is an intermediate step towards the sprout formation. Specifically, we imaged the progress of sprouting in the same areas of a live parental vessel at different time points of 1 hour, 3 hours, 7 hours and 28 hours of incubation in 100 ng/ml of VEGF (Fig. 4A-H). We found that all sprouts formed either directly from Stalks or from mini-sprouts, suggesting a non-observed transition from Stalk to mini-sprout due to observational timeframe limitations. Strikingly, however, not all mini-sprouts persisted and initiated sprout formation. Instead, many mini-sprouts retracted and new mini-sprouts formed during the time-course of the analysis. We then tracked a group of 118 cells that adopted the mini-sprout phenotype at least once over a period of 28 hours after VEGF exposure. Their state change dynamics was visualized using the Sankey diagram (Fig. 4I). The initial state of any sprouts or mini-sprouts was classified as the Stalk cell to reflect the hypothesized ‘salt-and-pepper ‘patterning structure, entirely consisting of either Tip or Stalk cells. When a mini-sprout retracted, it was newly marked as a Stalk cell. By the final time point of 28 hours of VEGF exposure, 45.8% of the cells that displayed the mini-sprout phenotype at least once during the experiment retracted back to the Stalk state, illustrating the highly dynamic phenotype of mini-sprout extensions and retractions. Of the remaining cells, 41.5% and 12.7% were classified to be either in the mini-sprout or sprout-leading Tip cell states, respectively. Although sprout formation continued throughout the experiment, the rate of conversion of mini-sprouts to full-fledged sprouts gradually decreased over time, with 13.6%, 2.9%, and 7.5 % of mini-sprouts becoming sprout-leading Tip cells in the time ranges of 1-3, 3-7 and 7-28 hours. respectively (Fig. 4K-M). In most cases (86.7%), sprouts emerged from *newly formed* mini-sprouts (i.e., the cells that were Stalk cells and then mini-sprouts in the preceding two time points), suggesting that mini-sprouts represent transient states rapidly converting to either the fully committed sprout state or to the Stalk state (Fig. 4 O). These observations raised the question of what might define the commitment of a mini-sprout to the sprout differentiation. We next addressed this question by analyzing the spatial distribution of fully formed sprouts over the observed area of the parental vessel.

**Figure 4.**
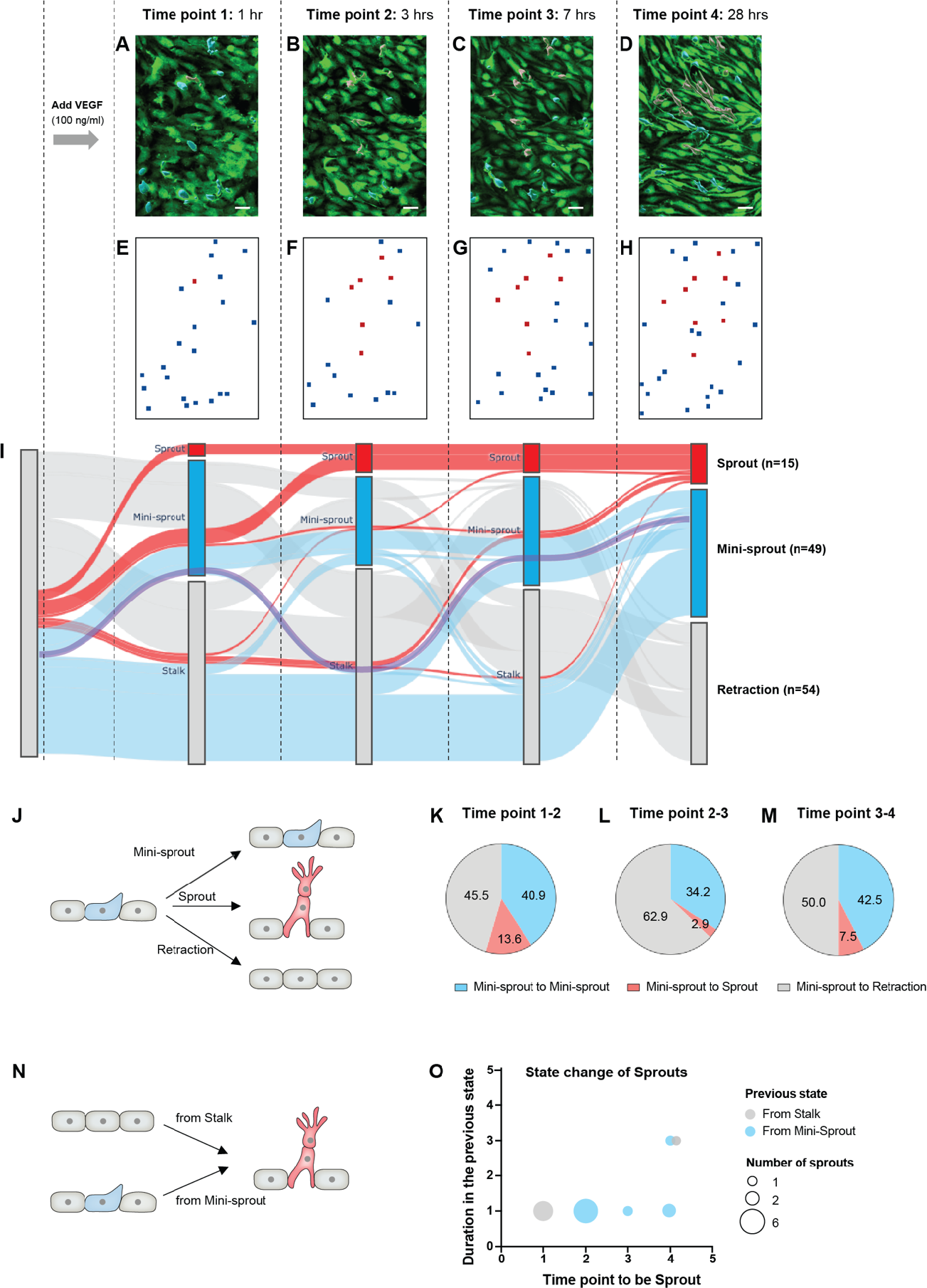
The dynamics of mini-sprout and sprout formation suggest frequent mini-sprout retractions, since only a subset of mini-sprouts becoming fully formed sprouts. GFP-expressing endothelium in the 3D vessel setup captured 1 hour (A), 3 hours (B), 7 hours (C), and 28 hours (D) after 100 ng/ml of VEGF treatment. Sprouts and mini-sprouts are identified by red and blue surface entities, respectively. Square marks representing the positions of sprouts (red) and mini-sprouts (blue) in the original images at each time point (E-H). (I) Sankey diagram demonstrating the dynamic state change of sprouts with red lines and mini-sprouts with blue lines throughout the time points. And grey lines represent mini-sprouts which ended up being retracted at the last observation, time point 4. A purple line shows an example of the state change from a Stalk (initially non-invading endothelial cell) to mini-sprout, retraction, mini-sprout, and mini-sprout at each time point. Only cells that that became mini-sprout at least once during the experiment are shown. (J) Different types of observed transitions between consecutive time points when starting from the mini-sprout state: maintain the mini-sprout state, become a sprout, or retract to the Stalk state. The ratio of states switched from mini-sprouts in the previous time point 1 (K), time point 2 (L), and time point 3 (M). (N) The two observed pathways to sprout formation between consecutive time points: direct Stalk to sprout or mini-sprout to sprout transition. Once a newly formed vessel becomes a sprout, it is permanently committed. (O) Duration of staying as a mini-sprout or a Stalk in the previous state before being committed to a sprout.

### Random uniform model accounts for spatial distribution of extending sprouts

While the NOTCH/VEGF mathematical model could quantitatively resolve Tip-Stalk spatial patterns, it did not capture the rate of cell fate switching or explained the commitment of Tip cells to lead the formation of a mature sprout. To identify the underlying principles of sprout initiation, we thus integrated the multicell model with several alternative phenomenological hypotheses (summarized in Fig. 5A-D). We then tested these hypotheses against the measured distributions of distances between sprouts and, in particular, the observation that sprouts were always separated by at least one non-sprout cell (blue bars in Fig. 5E). In these tests, we ensured that the results were normalized to the overall density of sprouts observed experimentally (see Method section: Phenomenological models of Sprout selection). The first straightforward hypothesis was that mini-sprouts commit to the sprout phenotype independently of the location of other forming sprouts, constituting the “cell-autonomous sprout selection” model (Fig. 5A-B). In this case, however, the corresponding model predicted mulTiple contacts between sprouts (black line in Fig. 5E), in sharp contrast with the experimental observation. The observation that most sprouts are in contact with at least another Tip (Supplementary figure 2A), but never in contact with another sprout, suggested a control mechanism where sprout selection inhibits nearby Tip cells from committing to the same fate. This led to two additional alternative hypotheses. In “repulsion between sprouts” model, it was assumed that sprouts cannot be in contact; therefore, Tip cells cannot commit to the sprout phenotype if already in contact with a sprout (Fig. 5C). In the “random uniform” model, it was assumed that sprouts are selected randomly, but maximizing their overall spread in the lattice (Fig. 5D; see Method section: Phenomenological models of Sprout selection). While both models correctly predicted sprouts to never be in contact, the “random uniform” model better described the cases where adjacent sprouts are separated by two or more cells (Fig. 5E).

**Figure 5.**
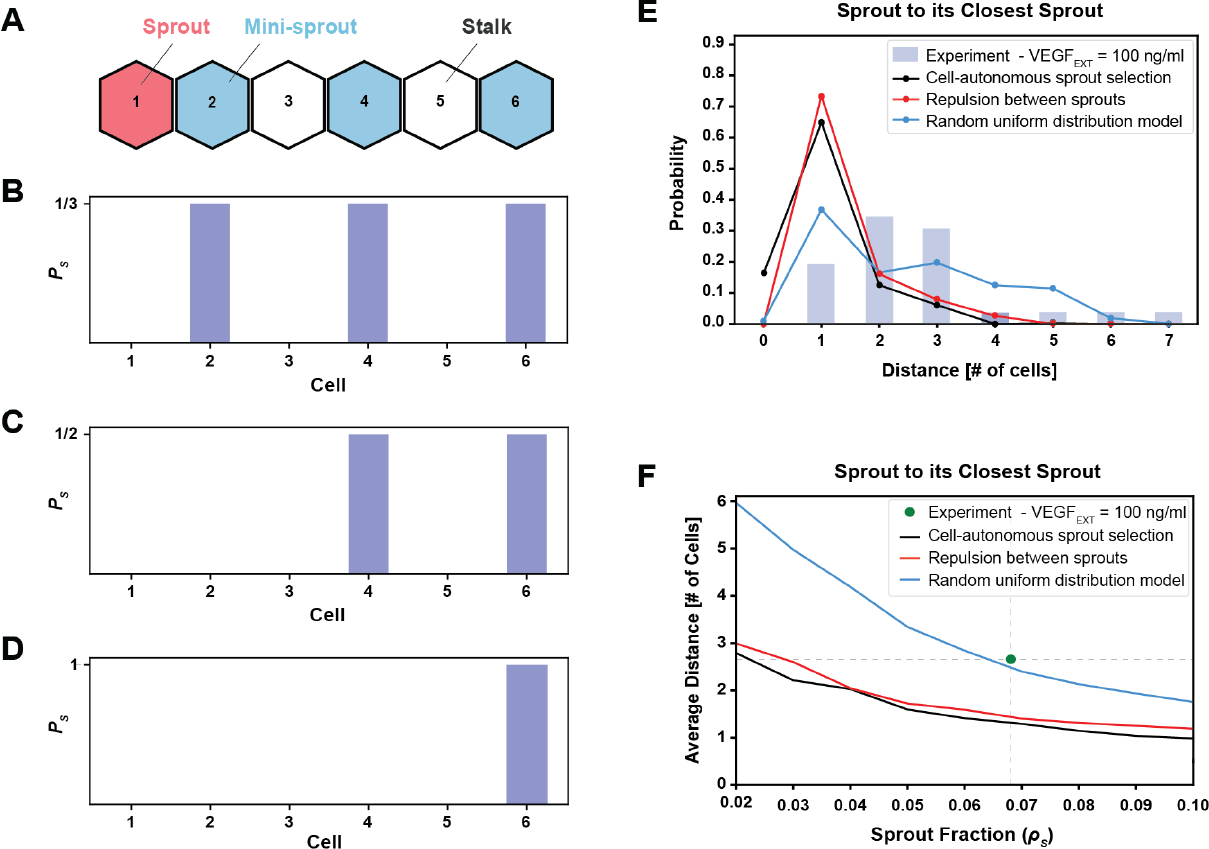
The phenomenological model favoring maximal sprout-sprout distances for a given number of sprouts (random uniform distribution) is most consistent with the experimental observations. (A) An example of 1-dimenstional Tip cell distribution, including a sprout and mini-sprouts, and Stalks pattern. (B) Sprout selection probability (*P*_*S*_) for the cell-autonomous model if a new sprout was added to the pattern of (A). Stalk cells cannot become sprouts, and existing mini-sprouts share the same selection probability. (C) Sprout selection probability (*P*_*S*_) for the sprout repulsion model. The leftmost mini-sprout cannot be selected because it is already in contact with an existing sprout, while the remaining two mini-sprouts share the same selection probability. (D) Sprout selection probability (*P*_*S*_) for the random uniform distribution model. The rightmost mini-sprout maximizes the distance to the existing sprout and is therefore the only viable selection. (E) Sprout distance distribution to its closest sprout neighbor. Blue bars indicate experimental results for VEGF_EXT_=100 ng/ml while black, red and blue lines depict the three different models of sprout selection (cell-autonomous, repulsion between sprouts, and random uniform distribution, respectively). (F) Average distance between a sprout and its closest sprout neighbor in the model as a function of sprout cell fraction in the lattice for the three proposed models of sprout selection. The green dot highlights the experimental sprout fraction and distance at VEGF_EXT_=100 ng/ml.

To test these mechanisms more rigorously, we computed the average distance (in cell numbers) between pairs of closest sprouts while also varying the number of sprouts allowed in the lattice (supplementary figure 2B-D), thus generating a curve of the typical sprout-sprout distancing as a function of sprout density in the lattice (Fig. 5F). The “random uniform” model predictions agreed very closely with the experimentally observed combination of sprout fraction and sprout-sprout distance, whereas the other two models greatly underestimated the distances between sprouts (Fig. 5F). Furthermore, while all models overestimated the fraction of the adjacent sprouts that are one cell away from the current sprout and underestimated the fractions of sprouts at greater distances, the deviation of the “random uniform” model predictions for this inter-sprout distribution was the lowest of all the models, again supporting the ‘random uniform’ model as the more likely to account for sprout selection (Fig. 5F).

### Fibronectin distribution may mediate sprout induction

Random distribution maximizing the distance between sprouts is similar to allelopathy models, accounting to spatial dispersion of species maximizing distance between them. In these models, the key postulated mechanism is inhibition of growth of individuals of the same species through a mutual suppression mechanism(16-18). Another analogous set of mechanisms are embedded within the concept of the Turing pattern formation, the key to which is diffusible negative feedback regulator setting spatial distribution of morphogenic features (19, 20). A variant of such mechanisms is a model postulating depletion of some ingredient that is key to the local growth, by its active redistribution towards the growing pattern features and depletion from the zones between them, rather than active mutual inhibition of the pattern forming units.

Given these prior models, we hypothesized that a similar mechanism may account for the dispersion patterns of sprouts. We focused on the extracellular matrix as a possible medium accounting for the positive and negative pattern-setting interactions. Indeed recently, it has been observed that collagen can be re-organized by the growing sprouts, so that it is concentrated around the extending sprouts and depleted elsewhere(21-23). We explored whether a similar distribution is also be observed for fibronectin, an ECM component that is critical for the formation of lumenized sprouts (24-27). Fibronectin expression levels in the vicinity of individual cells comprising the parental vessel and the emerging sprouts was experimentally assessed by immunostaining, with simultaneous cell identification using induced cytoplasmic GFP expression and nuclear staining, followed by 3D reconstruction (Figs. 6A-H). Untreated quiescent cells (No treatment) and mini-sprouts showed similar levels of fibronectin expression (Fig. 6H). Interestingly, the fibronectin expression was highly enriched at the base but not the Tip areas of the extending sprouts, suggesting that it may be a key determinant of sprout induction but not extension (Figs. 6B,C, &H).

**Figure 6.**
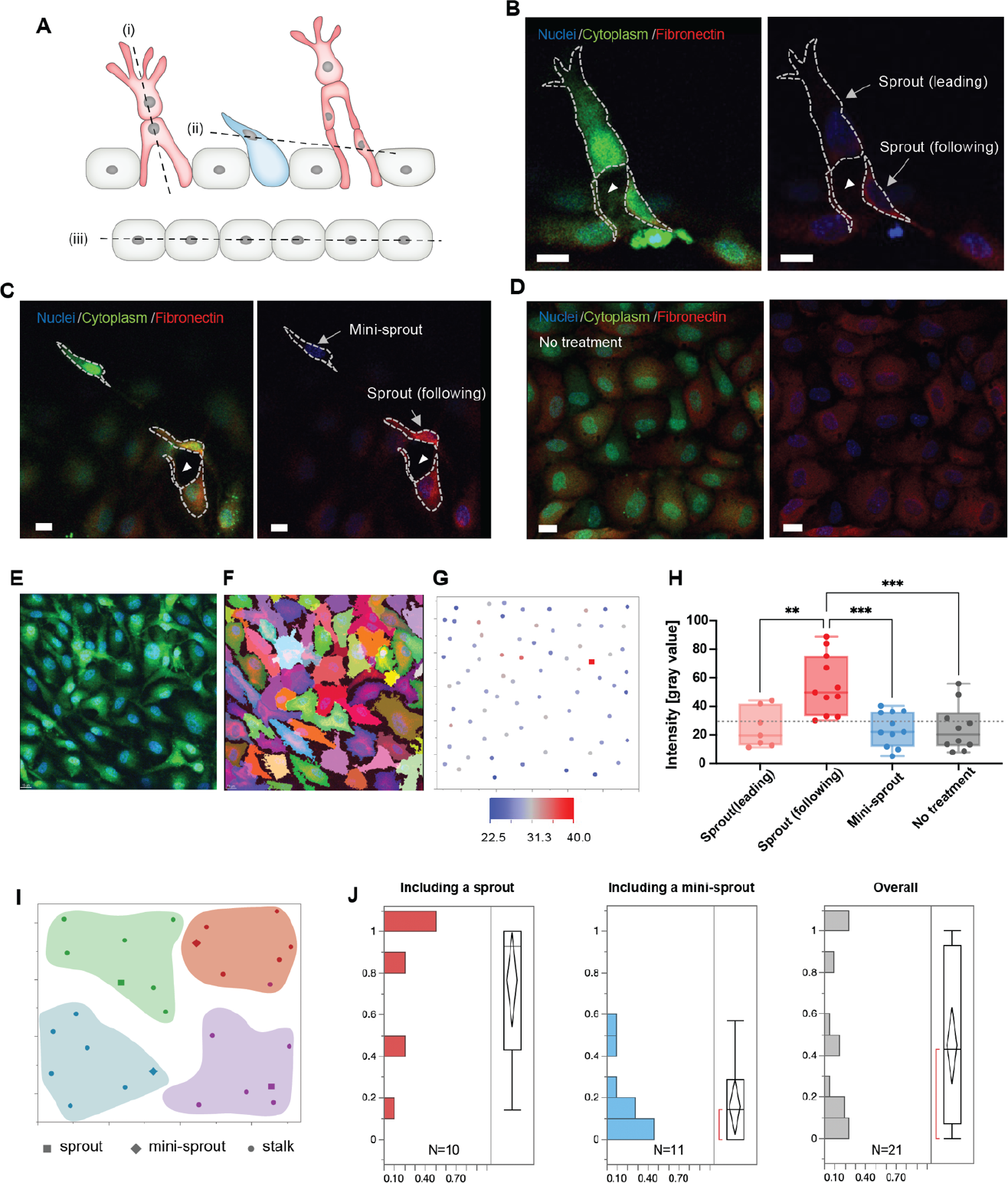
Fibronectin distribution on parental and newly formed vessels reveals preferred distribution at the bases of sprouts. (A) A schematic describing cross-sectional planes for subsequent confocal images: (i) for (B), (ii) for (C), and (iii) for (D). (B) Localization of fibronectin expression to a following cell than in a leading cell in a sprout. (C) Higher fibronectin expression in a sprout (a ‘following’ cell at the base of the sprout) than at a mini-sprout. (D) Intrinsic heterogeneity of fibronectin expression on quiescent endothelium displaying no mini-sprout or sprout formation. Images are 3D reconstructions of confocal z-stacks. Scale bars: 15 *μm*. Cells on the parental vessel were identified by GFP expression in the cytoplasm (E), then segmented (F). (G) Fibronectin intensity of each cell on the parental vessel is marked as a dot at the corresponding x and y positions of the cell centroids. Fibronectin intensities for sprouts (following cells) or mini-sprouts are indicated as squares. (H) Fibronectin intensity of leading cells and following cells of sprouts, mini-sprouts, and quiescent cells. (I) Cellular layer was segmented into groups containing 7 neighboring cells to assess the local environment for each group. (J) Distributions of the ratio of cells having fibronectin levels higher than a threshold, the minimum value of sprout in (H), in a group of 7 neighboring cells defined in (I), which included either a sprout or a mini-sprout. The overall distribution covers both regions.

We also examine the regional variation of fibronectin expression in larger areas, which was less variable and potentially more relevant to sprout extension. In particular, we accessed the ratio of cells having fibronectin levels higher than a threshold in groups of 7 cells around sprouts and mini-sprouts (Figs. 6I &J). The overall expression levels when all region types were combined was relatively uniform (Fig. 6J, third panel. Strikingly however, regions around sprouts (Fig. 6J, first panel) showed oppositely skewed patterns vs. mini-sprouts (Fig. 6J, second panel). Specifically, the fibronectin expression levels around sprouts were higher than the threshold, whereas the fibronectin levels around mini-sprouts were lower than the threshold. Altogether, these results supported the model in which fibronectin can indeed serve as a mediator of Turing-like induction of sprouting patterns, through re-modeling that enriches it at the points of sprout induction and depletes it at the points where Tip cells (mini-sprouts) are not stabilized to form full-fledged, lumenized sprouting bodies.

## Discussion

A major challenge of the analysis of tissue development and homeostasis is understanding of how differentiation into distinct cell types can be robustly achieved, while also being sensitive to various pro-differentiation and morphogenic cues and, potentially affected by molecular noise in biochemical reactions. In the context of angiogenesis, this challenge more specifically relates to enabling effective vascular morphogenesis through robust yet environmentally responsive differentiation of endothelial cells into Tip and Stalk cells states. This process is guided by the mulTiple pro-angiogenic cues, such as VEGF and by the local tissue organization and mediated by the paracrine Delta-NOTCH cell signaling. Recent mathematical models of this process(28, 29) have considered the robustness of the Delta-NOTCH differentiation to molecular noise, concluding that noisy biochemical reactions can both disrupt and enhance the spatial differentiation patterns, depending on the magnitude and spatial distribution (28). The effect of pro-angiogenic cues, such as VEGF on robustness of spatial differentiation patterning remains substantially less explored. Furthermore, compiling between modeling predictions and experimental validation has been challenged by the complexity of *in vivo* angiogenesis analysis, both on cellular and molecular levels.

Here, we integrated engineered experimental angiogenic assay and a spatially resolved computational modeling analysis to explore the spatially and temporally resolved effects of VEGF on angiogenic cell specification. Our results suggest that VEGF can have a dual role in inducing the initial Tip-Stalk cell differentiation. On the one hand, a low level of exogenous VEGF is essential to induce Delta-NOTCH signaling and the classical ordered ‘salt-and-pepper’ pattern, with approximately 25% of the cells adopting the Tip cell fate, as expected (30). This role of VEGF is conceptually similar to the classical case of EGF induced NOTCH signaling in *C. elegans* vulva development (31, 32). On the other hand, our results also suggest that an increase of VEGF levels can introduce disorder into this pattern, similar to order-disorder transition for various composite materials (12, 13), which may occur with increasing temperature and have the properties of a sharp phase transition. More specifically, high VEGF levels may play the role similar to an increased temperature in order-disorder transitions, leading to emergence of partially disordered ‘salt-and-pepper ‘structures. These disordered structures are characterized by higher than expected fractions of Tip cells and, consequently, an increased occurrences of otherwise disallowed adjacent Tip cells. Importantly, for all VEGF input levels and the resulting spatial patterns, the Tip-Stalk cell specification continued to be robust, although the degree of induction of NOTCH signaling was dependent on the VEGF dose. These results suggested that cell specification patterns deviating from the expected ‘salt-and-pepper’ one can develop not only due to noisy NOTCH signaling, as suggested by prior models, but also due to the control of the fractions of alternative cell states by the magnitude the pro-differentiation cue.

In spite of the observation that disordered ‘salt-and-pepper’ patterns still display robust differentiation, the spatial patterns of cell co-localization can generate inherent instabilities, e.g., due to two adjacent Tips mutually suppressing their fate selection through NOTCH signaling. Instabilities of this sort may increase the sensitivity of the differentiation process to additional cues and can also lead to facilitated dynamic switching of cell fates (e.g., Tip to Stalk and vice versa) over prolonged periods of time. Such instabilities might lead to oscillatory-like fluctuations of NOTCH signaling as observed in other differentiation processes (33-35) and in endothelial cell sheets under high VEGF concentration inputs(36). Importantly, these instabilities may also underlie the striking observation of the continuous retraction and extension of mini-sprouts (protruding Tip cells) observed in our study. This dynamic fate-switching behavior can thus represent a signature of an unstable differentiation process that may either stabilize, in response to additional cues, leading to a specific morphogenetic outcome, such as the extension of a stable sprout, or display a prolonged instability resulting in a lack of pronounced morphogenesis. Such poised but unstable states may be similar to the undifferentiated state of neurogenic progenitors displaying oscillatory NOTCH signaling, which can proceed to differentiation after the oscillation is resolved into a temporally stable NOTCH activity (37).

The cues stabilizing cellular differentiation and morphogenesis can vary and represent the signature of both global and local pro- and anti-angiogenic environments. For instance, our prior analysis indicated that exposure of model parental vessels to VEGF only rather than a more complete pro-angiogenic cocktail of various factors, can result in formation of mini-sprouts (and hence, effective Tip-Stalk cell differentiation), but not full-fledged sprouts (7). Sprout formation may also be modulated by the presence of mural cells and pro-inflammatory cytokines, which can indirectly modulate the NOTCH activity, but could also have additional effects, serving to stabilize a specific differentiation outcome. However, even if sprouts do form, it is not clear how their spatial distribution may arise and be potentially controlled by the environmental inputs. Our results argue that, for a given number of sprouts forming in the parental vessel, their mutual distances are maximized. This spatial distribution is consistent with a Turing-like mechanism (19, 20), implying the existence of a long-distance interaction inhibiting formation of new sprouts in the vicinity of the existing ones. Although the actual mechanism of putative Turning-like pattern formation is not fully elucidated here, our results are consistent with a variant of this regulatory behavior, in which a component of the extracellular matrix, fibronectin is actively redistributed by the nascent sprouts, with this ECM component being enriched at the points of sprout formation, but depleted in other zones, thus preventing sprout induction in the depleted areas. This redistribution is indeed equivalent to the classical Turing mechanisms that would involve generation of an explicitly inhibitory compound by the growing sprouts. Fibronectin has been implicated as a key pro-angiogenic ECM component, possibly due to its integration with VEGF, which further increases the plausibility of this mechanism (25). Interestingly, we find elevated levels of fibronectin at the bases of extending sprouts, but not at their Tips. This finding has two implications. First, fibronectin may be an important factor in the sprout induction rather than extension, hence the growing sprout can progress into the surrounding matrix beyond the area of enriched fibronectin, leaving it behind. Secondly, fibronectin can also promote lumen formation (26, 27), thus its enrichment at the bases of extending sprouts can further contribute to their lumenized structure. Furthermore, this mechanism can help explain the dramatic influence of cytokines, such as TNF, in preventing sprout formation, which can happen in sharp, TNF dose-dependent manner (7). Indeed, TNF avidly binds to fibronectin (24) and therefore, at high enough concentrations, would have a particular anti-angiogenic effect in the areas of increased fibronectin density, which according to the proposed mechanism would be the areas of incipient sprouts. This would dramatically increase its anti-sprouting effect, even if the effect of this cytokine on the NOTCH-dependent cell specification is more muted (7). Of note, another version of the Turing mechanism has recently been suggested to account for the branching of the sprouts, also involving formation of new Tip cells leading individual branches, although the molecular mechanism postulated in that analysis was distinct from the one proposed here (38, 39). These models, though plausible, will need to be further tested to ascertain causality of the proposed mechanisms, although at this stage the Turing-like mechanisms appear to be the best candidates to explain the experimental results we obtained.

Overall, our analysis supports the following dynamic view of angiogenic induction. The VEGF input can induce NOTCH signaling and formation of Tip cells that can behave as mini-sprouts. At lower VEGF input, a more ordered pattern of Tip cell induction can lead to formation of approximately 25% of Tip cells, but relatively few of these will become sprouts, reflecting lower sensitivity to additional ambient cues, promoting sprout formation. On the other hand, at higher VEGF inputs, a greater disorder of Tip cell patterning permits higher sensitivity to external cues, such as fibronectin, that can stabilize the sprouting. In addition, a more frequent co-localization of Tip cells under this condition enables initiation of a sprout from two adjacent cells, as observed in a fraction of cases in our experiments (the case in Table 1 involving formation of a sprout by two adjacent cells). The initial emergence of sprouts leads to a progressively less likely sprout formation and to the overall maximization of the distance between the sprouts. Both these observations are consistent with non-local inhibition of sprout formation around the sprouts that have already formed. A plausible mechanism for this inhibition and the overall pattern formation is the re-distribution of fibronectin from the zones between sprouts towards the incipient sprouts, constituting both a positive and negative feedback loops, commonly assumed in a variant of the Turing patterns from action. This mechanism is very sensitive to various local inputs distinct from VEGF and fibronectin, which can further influence the location and density of the developing blood vessels. These may include the effects of pericytes or local inflammatory environments, as mentioned above, but also other ECM components, such as collagens, that may enhance the protrusion of Tip cells and thus stabilize the emergent sprouts both directly and indirectly. Endothelial fate induction process may be interesting to contrast with other complex multicellular processes, including collective epithelial migration, where NOTCH signaling similarly modulates fine-grained patterns of leader and follower cells (40), and underscore the need in the future to develop more refined models that explicitly integrate the interconnections between biochemical and mechanical regulation of Tip-Stalk fate(41). The results in this study can further inform our understanding of angiogenesis in physiological and patho-physiological conditions. In particular, in many circumstances, the levels of VEGF is determined by the degree of hypoxia, which can be highly elevated following oxygen supply interruption, e.g., in wound healing or ischemia, or due to progression of neoplastic growth. Our results suggest that in these cases, formation of sprouts can be dysregulated due to higher incidences of co-localizations of prospective Tip cells. In addition, since these conditions are frequently accompanied by altered synthesis of ECM, the sprout density can increase, which may lead to formation of denser and less developed vascular beds frequently observed as a result of tumor angiogenesis(42, 43). Our results thus suggest that the disorder and higher plasticity of the endothelial cell fate speciation at higher VEGF inputs can be a key contributor to some pathological states associated with persistently hypoxic conditions.

The analysis presented here highlights the utility of combining experimental engineered vasculature models with mathematical analysis as integrated research platforms to gain a progressively better insight into angiogenesis in highly controlled micro-environments. Although, questions remain about the mechanistic underpinnings of the phenomena observed here and in related studies, the analysis enabled by these model systems can help formulate guiding hypotheses for further understanding of angiogenesis *in vivo*, while decoupling the complexities of the native angiogenic environments. We anticipate that the dynamic exploration presented here can help pave the way for further quantitative understanding of this key biological process.

## Materials and Methods

Chips for mimicking 3D angiogenesis *in vitro* were fabricated as introduced in our previous work(7). Fluorescence images of the *in vitro* model were acquired by Lecia scanning disk confocal microscopy, and the z-stack images were processed with IMARIS (Bitplane) to quantify Tip-Tip distance, to track the positions of sprouts and mini-sprouts, and to analyze fibronectin intensity distribution. We generalized existing mathematical models of the interconnected signaling between the NOTCH and VEGF pathways to a two-dimensional multicellular scenario (7, 44). The temporal dynamics of NOTCH (N), Delta (D), Jagged (J), NOTCH intracellular domain or NICD (I), and VEGF receptor (V_R_) in a cell were modeled with the parameter values presented in Table 2. The details of materials and methods can be found in *SI Appendix, Materials and Methods*.

**Table 2.** Parameter values for simulation. *Rescaled from previous model to match experimental observation.

| Parameter type | Parameter | Value | Units |
| --- | --- | --- | --- |
| Production | $N_0, D_0, J_0, V_{R0}$ | 1200, 1000, 800, 1000 | Molecule/hour |
| Degradation | $\gamma, \gamma_I$ | 0.1, 0.5 | 1/hour |
| Binding | $k_T, k_C$ | $2.5 \cdot 10^{-5}, 5 \cdot 10^{-4}$ | 1/(molecule hour) |
| Hill threshold | $I_0, V_0$ | 200, 80* | Molecules |
| Fold-change | $\lambda_N, \lambda_{I,D}, \lambda_{V,I}, \lambda_J, \lambda_V$ | 2, 0, 0, 2, 2 | Dimensionless |
| Hill coefficient | $n_N, n_{I,D}, n_{V,I}, n_J, n_V$ | 2, 2, 2, 5, 2 | Dimensionless |

## Acknowledgments

This work was supported by NIH through NCI U54 CA209992 (to A.L.); Mia Neri Foundation (to T.-Y.K.); The Center for Theoretical Biological Physics sponsored by the NSF (Grant PHY-2019745), NSF-PHY-2210291, CPRIT Scholar in Cancer Research sponsored by the Cancer Prevention and Research Institute of Texas (to J.N.O.); a Simons Foundation grant 594598 (to Q.N.). F.B. was supported by a National Science Foundation grant DMS1763272 (Q.N.).

## Notes

### Competing Interest Statement

The authors have declared no competing interest.

### Summary of Updates

The main article and supplementary information files have been revised based on reviewers' comments.

## References

1. R. H. Adams, K. Alitalo, Molecular regulation of angiogenesis and lymphangiogenesis. Nat Rev Mol Cell Biol 8, 464–478 (2007).

2. M. Potente, H. Gerhardt, P. Carmeliet, Basic and therapeutic aspects of angiogenesis. Cell 146, 873–887 (2011).

3. R. Benedito et al., The NOTCH ligands Dll4 and Jagged1 have opposing effects on angiogenesis. Cell 137, 1124–1135 (2009).

4. R. Blanco, H. Gerhardt, VEGF and NOTCH in Tip and Stalk cell selection. Cold Spring Harb Perspect Med 3, a006569 (2013).

5. W. Y. Wang, D. Lin, E. H. Jarman, W. J. Polacheck, B. M. Baker, Functional angiogenesis requires microenvironmental cues balancing endothelial cell migration and proliferation. Lab Chip 20, 1153–1166 (2020).

6. J. Liu et al., Synthetic extracellular matrices with tailored adhesiveness and degradability support lumen formation during angiogenic sprouting. Nat Commun 12, 3402 (2021).

7. T. Y. Kang et al., Pericytes enable effective angiogenesis in the presence of proinflammatory signals. Proc Natl Acad Sci U S A 116, 23551–23561 (2019).

8. M. B. Chen et al., On-chip human microvasculature assay for visualization and quantification of tumor cell extravasation dynamics. Nat Protoc 12, 865–880 (2017).

9. D. H. Nguyen et al., Biomimetic model to reconstitute angiogenic sprouting morphogenesis in vitro. Proc Natl Acad Sci U S A 110, 6712–6717 (2013).

10. M. Boareto et al., Jagged-Delta asymmetry in NOTCH signaling can give rise to a Sender/Receiver hybrid phenotype. Proc Natl Acad Sci U S A 112, E402–409 (2015).

11. F. Bocci, J. N. Onuchic, M. K. Jolly, Understanding the Principles of Pattern Formation Driven by NOTCH Signaling by Integrating Experiments and Theoretical Models. Front Physiol 11, 929 (2020).

12. Y. Yang et al., Deciphering chemical order/disorder and material properties at the single-atom level. Nature 542, 75–79 (2017).

13. K. Bu et al., Nested order-disorder framework containing a crystalline matrix with self-filled amorphous-like innards. Nat Commun 13, 4650 (2022).

14. O. Shaya et al., Cell-Cell Contact Area Affects NOTCH Signaling and NOTCH-Dependent Patterning. Dev Cell 40, 505–511 e506 (2017).

15. M. Kwak et al., Adherens junctions organize size-selective proteolytic hotspots critical for NOTCH signalling. Nat Cell Biol 24, 1739–1753 (2022).

16. D. L. Liu, M. An, I. R. Johnson, J. V. Lovett, Mathematical Modeling of Allelopathy. III. A Model for Curve-Fitting Allelochemical Dose Responses. Nonlinearity Biol Toxicol Med 1, 37–50 (2003).

17. G. S. Fraenkel, The raison d’etre of secondary plant substances; these odd chemicals arose as a means of protecting plants from insects and now guide insects to food. Science 129, 1466–1470 (1959).

18. R. J. Willis, The history of allelopathy (Springer, Dordrecht, 2007), pp. xiv, 316 p. : ill., facsims., map, ports.

19. A. M. Turing, The chemical basis of morphogenesis. 1953. Bull Math Biol 52, 153–197; discussion 119-152 (1990).

20. P. K. Maini, R. E. Baker, C. M. Chuong, Developmental biology. The Turing model comes of molecular age. Science 314, 1397–1398 (2006).

21. X. Feng, M. G. Tonnesen, S. A. Mousa, R. A. Clark, Fibrin and collagen differentially but synergistically regulate sprout angiogenesis of human dermal microvascular endothelial cells in 3-dimensional matrix. Int J Cell Biol 2013, 231279 (2013).

22. A. Senk, V. Djonov, Collagen fibers provide guidance cues for capillary regrowth during regenerative angiogenesis in zebrafish. Sci Rep 11, 19520 (2021).

23. N. D. Kirkpatrick, S. Andreou, J. B. Hoying, U. Utzinger, Live imaging of collagen remodeling during angiogenesis. Am J Physiol Heart Circ Physiol 292, H3198–3206 (2007).

24. R. Alon et al., TNF-alpha binds to the N-terminal domain of fibronectin and augments the beta 1-integrin-mediated adhesion of CD4+ T lymphocytes to the glycoprotein. J Immunol 152, 1304–1313 (1994).

25. E. S. Wijelath et al., Heparin-II domain of fibronectin is a vascular endothelial growth factor-binding domain: enhancement of VEGF biological activity by a singular growth factor/matrix protein synergism. Circ Res 99, 853–860 (2006).

26. S. Astrof, R. O. Hynes, Fibronectins in vascular morphogenesis. Angiogenesis 12, 165–175 (2009).

27. K. J. Bayless, R. Salazar, G. E. Davis, RGD-dependent vacuolation and lumen formation observed during endothelial cell morphogenesis in three-dimensional fibrin matrices involves the alpha(v)beta(3) and alpha(5)beta(1) integrins. Am J Pathol 156, 1673–1683 (2000).

28. M. Galbraith, F. Bocci, J. N. Onuchic, Stochastic fluctuations promote ordered pattern formation of cells in the NOTCH-Delta signaling pathway. PLoS Comput Biol 18, e1010306 (2022).

29. Y. L. Koon, S. Zhang, M. B. Rahmat, C. G. Koh, K. H. Chiam, Enhanced Delta-NOTCH Lateral Inhibition Model Incorporating Intracellular NOTCH Heterogeneity and Tension-Dependent Rate of Delta-NOTCH Binding that Reproduces Sprouting Angiogenesis Patterns. Sci Rep 8, 9519 (2018).

30. J. R. Collier, N. A. Monk, P. K. Maini, J. H. Lewis, Pattern formation by lateral inhibition with feedback: a mathematical model of delta-NOTCH intercellular signalling. J Theor Biol 183, 429–446 (1996).

31. H. Shin, D. J. Reiner, The Signaling Network Controlling C. elegans Vulval Cell Fate Patterning. J Dev Biol 6 (2018).

32. A. S. Yoo, C. Bais, I. Greenwald, CrosStalk between the EGFR and LIN-12/NOTCH pathways in C. elegans vulval development. Science 303, 663–666 (2004).

33. Y. Zhang et al., Oscillations of Delta-like1 regulate the balance between differentiation and maintenance of muscle stem cells. Nat Commun 12, 1318 (2021).

34. O. F. Venzin, A. C. Oates, What are you synching about? Emerging complexity of NOTCH signaling in the segmentation clock. Dev Biol 460, 40–54 (2020).

35. R. Kageyama, Y. Niwa, H. Shimojo, T. Kobayashi, T. Ohtsuka, Ultradian oscillations in NOTCH signaling regulate dynamic biological events. Curr Top Dev Biol 92, 311–331 (2010).

36. B. Ubezio et al., Synchronization of endothelial Dll4-NOTCH dynamics switch blood vessels from branching to expansion. Elife 5 (2016).

37. H. Shimojo, T. Ohtsuka, R. Kageyama, Oscillations in NOTCH signaling regulate maintenance of neural progenitors. Neuron 58, 52–64 (2008).

38. S. Guo, M. Z. Sun, X. Zhao, Wavelength of a Turing-type mechanism regulates the morphogenesis of meshwork patterns. Sci Rep 11, 4813 (2021).

39. H. Xu, M. Sun, X. Zhao, Turing mechanism underlying a branching model for lung morphogenesis. PLoS One 12, e0174946 (2017).

40. S. A. Vilchez Mercedes et al., Decoding leader cells in collective cancer invasion. Nat Rev Cancer 21, 592–604 (2021).

41. O. Stassen, T. Ristori, C. M. Sahlgren, NOTCH in mechanotransduction - from molecular mechanosensitivity to tissue mechanostasis. J Cell Sci 133 (2020).

42. E. Ruoslahti, Specialization of tumour vasculature. Nat Rev Cancer 2, 83–90 (2002).

43. A. S. Chung, J. Lee, N. Ferrara, Targeting the tumour vasculature: insights from physiological angiogenesis. Nat Rev Cancer 10, 505–514 (2010).

44. M. Boareto, M. K. Jolly, E. Ben-Jacob, J. N. Onuchic, Jagged mediates differences in normal and tumor angiogenesis by affecting Tip-Stalk fate decision. Proc Natl Acad Sci U S A 112, E3836–3844 (2015).

