## Supplementary Information for "Spatial-temporal order-disorder transition in angiogenic NOTCH signaling controls cell fate specification"

**This PDF file includes:**

Supplementary Materials and Methods

Figures S1 to S2

Supplementary References

### Supplementary Materials and Methods

**Fabrication of 3D vessel chip**
Chips for mimicking 3D angiogenesis *in vitro* were fabricated as introduced in our previous work(1). The chip for live-cell imaging consists of a PDMS chamber, an engineered blood vessel embedded in collagen gel, and the whole construct was placed on a glass-bottom dish (Fig. 1A). A single line mold (D: 200 -250 um, L: 10 mm) of PLA which has a semicircular cross-section was deposited on a petri dish with a 3D printer (Ultimaker). After a PDMS chamber was put on the mold, pre-mixed collagen solution (5mg/ml, Type 1 collagen, BD) according to manufacturer's protocol was injected through a hole on the PDMS chamber and collagen polymerization was induced on ice for 30 min and at 37 ℃ for 1.5 hours. The PDMS chamber including the collagen construct was carefully peeled off, then the bottom side was sealed with a glass bottom of a dish. Laminin solution (60ug/ml in PBS, Sigma Aldrich) was injected into the channel pre-made in collagen, then the chip was flipped upside down and incubated at 37 ℃ for 1 hour. Finally, endothelial cell suspension ($1\times10^7$cells/ml) was injected into the channel, then the chip was flipped upside down again and incubated at 37 ℃ for 1 hour to allow cells to attach to the lumen. Once endothelial cells form a confluent monolayer on the channel surface, a fresh medium was injected into the engineered blood vessel and changed every 12 hours before the treatment with pro-angiogenic factors.

**Cell culture and induction of angiogenesis**
GFP-expressing human brain endothelial cells were purchased from Angio-Protemie and cultured in a growth medium of M199 (Gibco) supplemented with 20% FBS (Life Technologies), 1% HEPES (Thermo Scientific), 1% Glutamax (Thermo Fisher), 1% antibiotic-antimycotic (Thermo Fisher), Heparin (25mg/500ml, Sigma Aldrich), and endothelial cell growth supplement (Sigma Aldrich). For inducing angiogenesis in the 3D vessel chip, the growth medium was supplemented with 40ng/ml of basic fibroblast growth factor (bFGF, Thermo Fisher), 500nM of Sphingosine-1-phosphate (S1P, Sigma Aldrich), and 75ng/ml of phorbol myristate acetate (PMA, Sigma Aldrich). And 100 ng/ml or 10 ng/ml of VEGF was added on the top reservoir of the 3D vessel chip to create a VEGF gradient toward the engineered blood vessel (Fig. 1B).

**Mathematical model of VEGF/NOTCH signaling**
We generalize existing mathematical models of the interconnected signaling between the NOTCH and VEGF pathways to a two-dimensional multicellular scenario (1, 2). The temporal dynamics of NOTCH (N), Delta (D), Jagged (J), NOTCH intracellular domain or NICD (I), and VEGF receptor ($V_R$) in a cell are modeled with ordinary differential equations:

$$\frac{dN}{dt} = N_0H^+(I) - k_TN(D_{EXT} + J_{EXT}) - k_CN(D + J) - \gamma N \quad (1a)$$
$$\frac{dD}{dt} = D_0H^-(I)H^+(V_RV_{EXT}) - k_TN_{EXT}D - k_CND - \gamma D \quad (1b)$$
$$\frac{dJ}{dt} = J_0H^+(I) - k_TN_{EXT}J - k_CNJ - \gamma J \quad (1c)$$
$$\frac{dI}{dt} = k_TN(D_{EXT} + J_{EXT}) - \gamma_I I \quad (1d)$$
$$\frac{dV_R}{dt} = V_{R0}H^-(I) - k_TV_RV_{EXT} - \gamma V_R \quad (1e)$$

NOTCH, Delta, Jagged, and VEGF receptor are produced with basal rates $N_0, D_0, J_0, V_{R0}$ and are degraded with rate constant $\gamma$. The basal production rates are modulated by NICD that transcriptionally activates NOTCH and Jagged while inhibiting Delta and VEGFR via shifted Hill functions:

$$H^S(I, I_0, n, \lambda) = \frac{1 + \lambda\left(\frac{I}{I_0}\right)^n}{1 + \left(\frac{I}{I_0}\right)^n} \quad (2)$$

Where $I_0$ is a threshold NICD level, $n$ is the Hill coefficient, and $\lambda$ represents the target's production rate fold-change at high NICD concentrations ($I \gg I_0$). Therefore, $\lambda > 1$ implies transcriptional activation while

$\lambda < 1$ implies transcriptional inhibition. For brevity, activating and inhibiting Hill functions are denoted by $H^{+}(I)$, $H^{-}(I)$, respectively.
Receptors and ligands can bind to external ligands/receptors with binding rate constant $k_T$. In the case of NOTCH signaling, $N_{EXT}$, $D_{EXT}$, and $J_{EXT}$ represent the average levels of NOTCH, Delta and Jagged in the six nearest neighbor cells on the hexagonal lattice. Conversely, VEGF is modeled as an external signal provided to all endothelial cells; therefore, all cells are exposed to the same fixed level ($V_{EXT}$). Moreover, NOTCH receptors and ligands can bind within the same cell with a rate constant $k_C$, which results in the degradation of the ligand-receptor complex without any downstream signaling (cis-inhibition). NICD is released upon binding of NOTCH receptors with external ligands and degraded with rate constant $\gamma_I$. Finally, VEGF receptors can bind to external VEGF ligands, thus creating activated VEGF receptor ($V_R V_{EXT}$), which in turn inhibits the production of Delta. Details on parameter values are presented in Table 2. Compared to previous models, we rescaled the Hill function threshold for VEGF-mediated activation of Delta in order to match quantitatively the ratio of Tip/Stalk cells between model and experiment when the external input is $V_{EXT} = 100$ ng/ml.
In this project, we focus specifically on the regulation of VEGF on the NOTCH-Delta signaling pathway (as shown in Fig. 2A). A strong NOTCH-Jagged interaction can suppress Tip-Stalk differentiation and instead lead to a hybrid Tip/Stalk phenotype, which is beyond the scope of the current study. For completeness, we maintained the integrity of the entire circuit structure including NOTCH-Jagged interactions (eq. 1c) but fixed the production rate of Jagged at a low level that does not interfere with the bistable behavior of the VEGF/NOTCH circuit.

**Definition of Tip cells in the model**
Tip cells – usually defined as high Delta, low NOTCH cells – are defined in this model as cells with Delta levels larger than $10^3$ molecules. This definition is motivated by observing that the distribution of cellular Delta at steady state in the multicell model is always bimodal irrespectively of the level of external VEGF input (see figure 2B), with a large separation between the population of cells with low Delta levels (the Stalk cells) and the population of cells with high Delta levels (the Tip cells). This phenomenological definition suffices here due to the deterministic nature of the model; otherwise, more complex approaches would be necessary in presence of stochastic fluctuations.

**Simulation details**
All results are calculated on a 30x30 hexagonal lattice with periodic boundary conditions. Initially, the lattice is prepared with randomized initial conditions where the initial levels of N, D, J, I, and $V_R$ within each cell are sampled from uniform distributions. Afterwards, the lattice equilibrates for 100 hours without any VEGF input to simulate the seeding of endothelial cells before VEGF is provided.

**Quantification of Tip-Tip distance in experiments**
Fluorescence microscopy images of DAPI/phalloidin staining from our previous work(1), two images for each VEGF condition, were used to quantify the distance between Tips. Before counting Tip-Tip distance, images were pre-processed to clean up intricate 3D images as depicted in Figure 3A with IMARIS(Bitplane). First, all newly formed vessels were separated from parental vessels, identified as surface entities, and labeled with different colors depending on lumen formation: red for sprout and blue for mini-sprout. And all nuclei were detected as spots (Fig. 1A, Label nuclei on the parental vessel and Tip region). In this step, nuclei on the parental vessel region were marked as Stalks and the nuclei in sprout or mini-sprout surfaces were marked as Tips. Second, the nuclei of Stalks on the parental vessel region were re-examined (Fig. 3A, Re-label Tip and Stalk nuclei). If they are connected to sprout or mini-sprout surfaces (Fig. 3A (1) & (2)), they were re-labeled as nuclei of sprouts or mini-sprouts and all other nuclei in the sprout or the mini-sprout were deleted. If a nucleus of a sprout or a mini-sprout is not in the parental vessel, the closest nucleus to the parental vessel was labeled as a sprout or a mini-sprout and all other nuclei were deleted (Fig. 3A (3)). By going through these steps, we could derive 2D conceptual Tip-Stalk arrangements from 3D images, which are comparable to 2D Tip-Stalk patterns from mathematical modeling. The pre-process is based on our assumption that the Stalk cells found in the extending sprouts emerge *de novo* through cell proliferation, rather by Stalk cell migration from the parental vessel.
In this study, we defined the distance between two Tips as the number of cells measured in 'cell hops', i.e., the minimal number of intermediate cells between randomly chosen pairs of Tip cells. For example, in Figure 3B, the distances from Tip 1 to Tip 2 and to Tip 3 are 0 and 1, respectively. From Tip 1 to Tip 4, the distance

is 3 in the minimal cell hops marked with gray arrows which pass through another Tip. Another cell hops marked with red arrows does not include any Tips, but the distance is 4. In this case, we discarded Tip 4 in the quantification of Tip-Tip distance from Tip 1. The same rule was applied in analyzing Tip-Tip distance from mathematical modeling (Figure 3C).

**Quantification of Tip-Tip distance in modeling**

Upon complete lattice equilibration, cells naturally separate into two distinct groups based on low or high expression of Delta irrespectively of the external VEGF input (see Fig. 2B). Therefore, high-Delta cells are labeled as Tips and low-Delta cells are labeled as Stalk. The algorithm to compute distances between Tip cells on the hexagonal lattice follows the following two steps: (1) Computing the shortest path between a given pair of cells on the hexagonal grid; and (2) filtering out the measurement if the cell pair is 'shielded' by another Tip cell. First, the shortest path is defined as the minimum number of intermediate cells that connect the two cells of interest. Considering two cells with coordinates $(x_1, y_1)$ and $(x_2, y_2)$, their distance has different values based on the cells relative position. In the simple case where $dx = 0$ or $dy = 0$, the distance is $d = dx + dy$. Else, if the distance is $d = dx + dy - 1$ (1) if $dy$ is even of (2) if $dy$ is odd but $y_1$ is even. In any other case, the distance is $d = dx + dy$. Next, to maintain consistency with the experimental statistics, the distance is not included in the Tip cell distribution if there is at least one intermediate Tip cell 'shielding' the two Tip cells of interest. In the experimental protocol, an intermediate cell 'shields' the two Tip cells if the straight line connecting the centers of the two cells passes through the shielding cell. Therefore, we search the intercept between the line connecting the pair of Tip cells and a circle centered at the position of the shielding cell with radius of 0.5. Assuming a circular geometry for the shielding cell removes artifacts emerging due to the polygonal shape of cells in the hexagonal lattice. The Sprout-Sprout distance statistics presented in Fig. 5E are similarly computed by applying the same algorithm to the set of Sprout cells in the lattice (see the following section for details of Sprout selection).

**Phenomenological models of Sprout selection**

We developed three separate models of sprout selection: "Cell-autonomous", "Repulsion", and "Random uniform distribution". For all three cases, cells equilibrate in the hexagonal lattice. Then, Tip cells are defined based on high-Delta expression. This analysis produces a discrete snapshot of Tip and Stalk cells on the hexagonal lattice. When comparing the model prediction to the 100ng/ml VEGF dosage experiment, we constrain the Sprout selection models to reproduce the experimental Sprout fraction. In the "Cell-autonomous" model, a fraction of the Tip cells is randomly selected and defined as Sprouts independently from the phenotype of their neighbors. In the "Repulsion" model, a fraction of Tip cells is randomly selected as Sprouts with the additional constraint that Sprouts cannot be in direct contact. To implement this constraint, we iteratively select Tips one at a time. If the selected Tip is not already in contact with a previously selected Sprout, it is promoted to the Sprout state; otherwise, a new Tip cell is selected. The iteration stops when the target number of Sprouts is reached. Finally, in the "Random uniform distribution" model, Tips are selected as Sprouts to maximize their overall spread in the lattice. To implement this constraint, we first select a random Tip and promote it to the Sprout state. Then, the furthest Tip from the newly selected Sprout is selected and promoted to the Sprout state as well. Afterwards, a max-distance function is defined as the sum of pairwise distances between all Sprouts already selected in the lattice. At each following iteration, the Tip cell that maximizes the distance function is selected as a Sprout. The iteration stops when the target number of Sprouts is reached.

**Sprout/mini-sprout tracking and analysis**

3D vessel chips were kept in a $CO_2$ incubator at 37 °C and taken out only for imaging at the timepoint depicted in Figures 4A-D. GFP-expressing endothelial cells on a chip at the same position were imaged with a 20x water-immersion objective attached to Lecia scanning disk confocal microscope. IMARIS (Bitplane) was used to separate all newly formed vessels from the parental vessel and to label them with different colors depending on lumen formation: sprout and mini-sprout. The positions of sprouts and mini-sprouts were simply marked with red and blue squares, respectively (Figs. 4E-H). Then the appearance, disappearance, or the transition from blue to red squares were recorded and tracked throughout the time points. Data from two independent samples were combined to plot the Sankey diagram demonstrating the dynamic change of sprouts and mini-sprouts (Fig. 4I). The analysis was performed using the Python graphing library Plotly.

**Immunofluorescence staining of Fibronectin and analysis**

All reagents for fibronectin immunostaining were injected through the vessel on a chip and kept as follows: 4% (wt/wt) formaldehyde for 1 hour, 0.1% triton-X for 1hour, 10% goat serum for 1hour, fibronectin antibody (Abcam) at 1:50 dilution for 1 day, secondary antibody (anti-rabbit 594 Thermo Scientific) at 1:100 dilution with Hoechst at 1/200 dilution for 1 day. 3D volumes at 10 positions from two independent samples of VEGF 100 ng/ml treatment and 2 positions from control (No treatment) were imaged with a 20x water-immersion objective attached to a Lecia scanning disk confocal microscope. Cells on the parental vessel were identified by GFP expression in the cytoplasm (Fig. 6E), then segmented (Fig. 6F) with IMARIS (Bitplane). The averaged fibronectin intensity value of each 3D cell volume and its position were acquired in IMARIS, then each cell was marked as a circle at the corresponding x and y positions (Fig. 6G). The averaged fibronectin intensity values and positions of sprouts and mini-sprouts were acquired by identifying cells as surface entities in IMARIS. The averaged fibronectin intensity values of sprouts (following cell) and mini-sprouts were marked as squares at the corresponding x and y positions (Fig. 6G). 10 sprouts, 11 mini-sprouts induced by VEGF 100 ng/ml, and 10 quiescent cells without VEGF treatment were selected to compare fibronectin intensity values (Fig. 6H). Sprouts often had two compartments of a leading cell and following cells where lumenization occurs. We distinguished the leading sprout and the following sprout for the fibronectin intensity comparison (Fig. 6H). If a sprout was composed of a single cell with lumen, the cell was classified as the following cell. 7 neighboring cells including a sprout or a mini-sprout in each image were grouped (Fig. 6I) to analyze the distribution of the ratio of cells having fibronectin levels higher than a threshold value of 30 (dashed line in Fig. 6H). From 10 images, 21 groups were acquired and used for the analysis (Fig. 6J).

**Statistical analysis**

One-way ANOVA was performed in GraphPad Prism 9.0.0 followed by Tukey's mulTiple comparisons test. Differences between pairs were concluded to be significant if they had adjusted p values less than 0.05.

#### Supplementary Figures

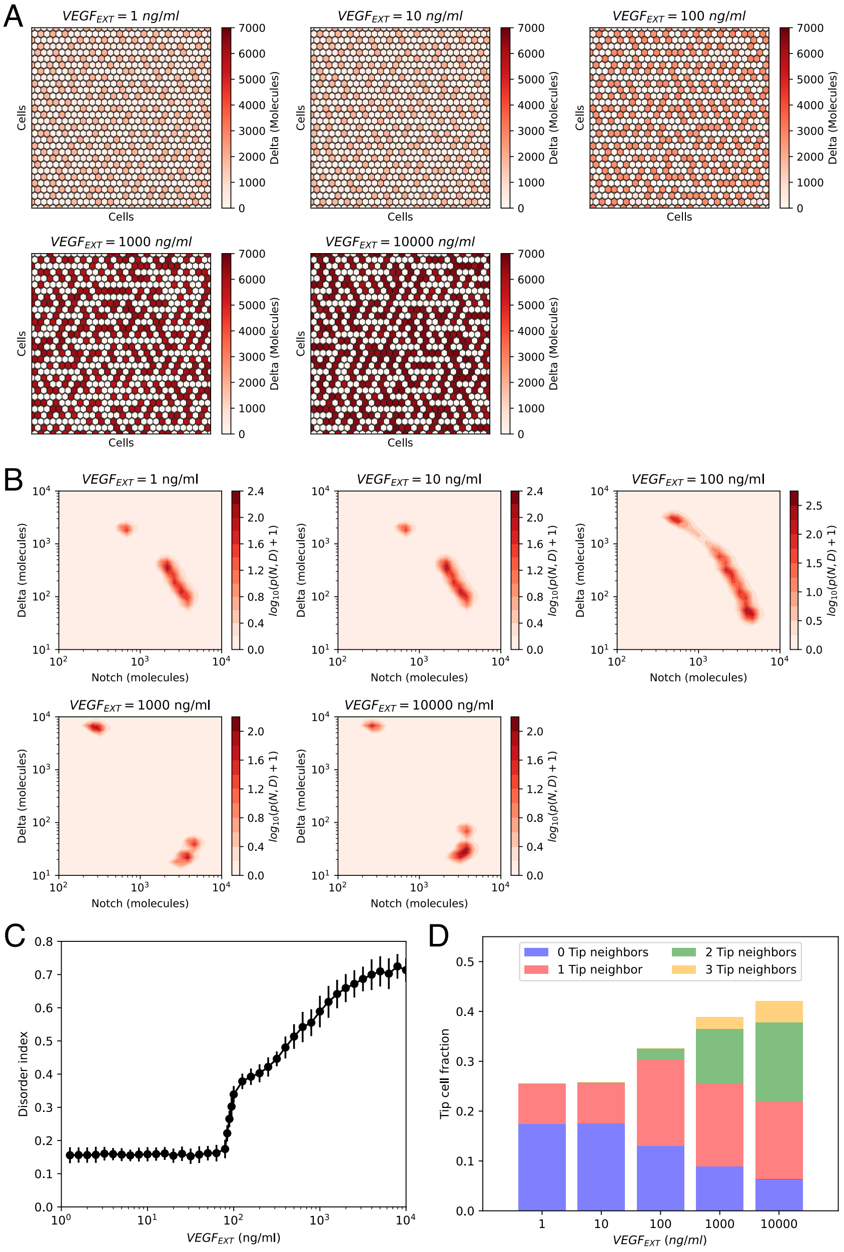


**Fig. S1. Order-Disorder transition in the NOTCH-Delta-VEGF multicell model. (A)** Steady state patterns for increasing levels of VEGF signal. Red heatmaps show the levels of Delta in each lattice cell. **(B)** Pseudopotential landscape of the joint (NOTCH, Delta) intracellular levels for increasing levels of VEGF signal. **(C)** The disorder index of the pattern as a function of external VEGF signal. The disorder index is defined as the fraction of "incorrect" Tip-Tip contacts in the lattice. **(D)** Statistics of Tip-Tip contacts as a function of external VEGF signal. For each VEGF input level on the x-axis, the bar height represents the overall fraction of Tip cells in the pattern, while the color-coding classifies Tip cells based on the number of "incorrect" Tip nearest neighbors. For panels (B-C-D), results are averaged over n=50 independent simulations starting from randomized initial conditions on a 30x30 hexagonal lattice with periodic boundary conditions (see methods for details).

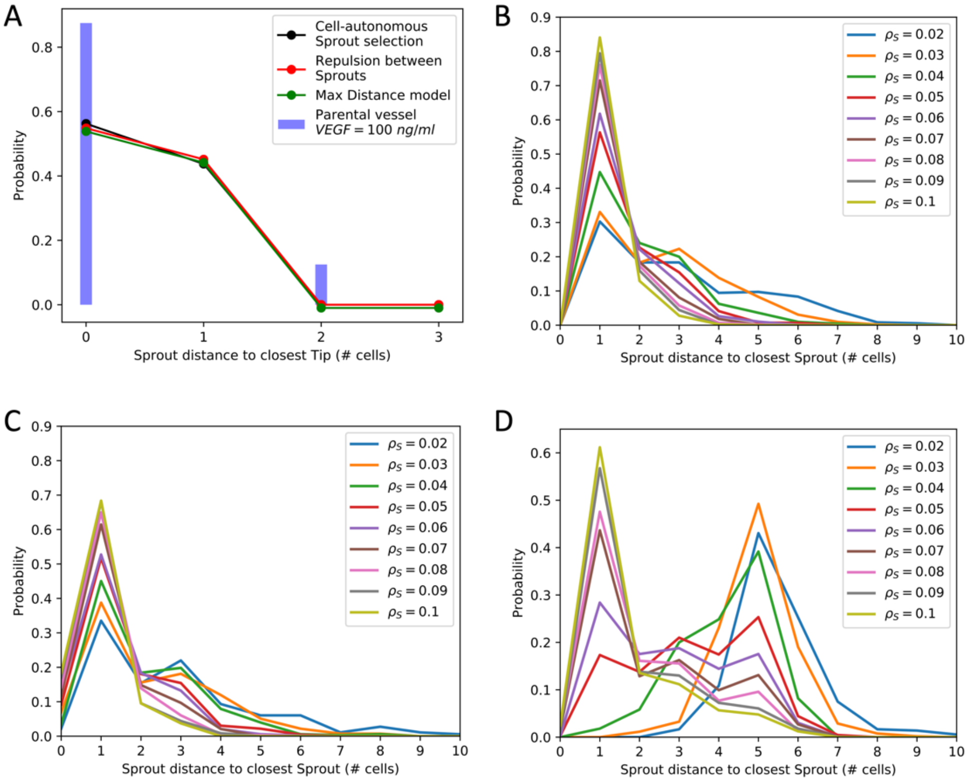


**Fig. S2. Phenomenological models of Sprout selection. (A)** Detailed distribution of Sprout distance to its closest Tip neighbor. Solid blue bars depict experimental measurements for the VEGF=100 ng/ml experiment, while black, red, and green lines depict the predicted distance distributions for the cell-autonomous, repulsion, and random uniform Sprout selection models. **(B-C-D)** Distribution of Sprout distance to its closest Sprout neighbor predicted by the model. Different curves indicate the Sprout distance distribution for models with increasing Sprout cell density ranging from 2% to 10% (the experimental sprout density at VEGF=100 ng/ml is $\rho_S = 7\%$). Panels B, C, and D depict predictions for the cell-autonomous, repulsion between sprouts, and random uniform sprout selection models, respectively. In all panels, the VEGF level is fixed to VEGF=100 ng/ml to match the experimental model.

### Supplementary References


1. T. Y. Kang *et al.*, Pericytes enable effective angiogenesis in the presence of proinflammatory signals. *Proc Natl Acad Sci U S A* **116**, 23551-23561 (2019).
2. M. Boareto, M. K. Jolly, E. Ben-Jacob, J. N. Onuchic, Jagged mediates differences in normal and tumor angiogenesis by affecting Tip-Stalk fate decision. *Proc Natl Acad Sci U S A* **112**, E3836-3844 (2015).
